# Tolerance toward foreigners in ants requires chronic exposure for establishment but only sporadic exposure for maintenance

**DOI:** 10.64898/2025.12.10.693507

**Authors:** Tiphaine P. M. Bailly, Matteo Rossi, Stephany Valdés-Rodríguez, Thomas Schmitt, Erik T. Frank, Daniel J. C. Kronauer

## Abstract

Social insects discriminate between foreigners and members of their own colony via complex olfactory cues. How plastic this discriminatory behavior is, and whether and under what circumstances ants can learn to accept genetically distinct individuals as nestmates, is poorly understood. Here, we study this question in the clonal raider ant, *Ooceraea biroi*, which provides unparalleled experimental control over the genotype of individuals and the genotypic composition of colonies. Using a cross-fostering design with mixed-genotype colonies of wild-type and transgenically labelled individuals, we show that ants become non-aggressive specifically toward their foster genotype. This tolerance is transient, and aggression resumes after two weeks of being isolated from the foster colony. However, even sporadic re-exposure to individuals from the foster colony is sufficient to maintain tolerance for over a month, while the same paradigm fails to establish tolerance in the first place. This shows that non-nestmate discrimination is remarkably plastic and that, once established, tolerance toward foreigners can be maintained by only intermittent contact. These dynamics echo general principles of social learning and contact-dependent tolerance described in other social species, including humans.

## INTRODUCTION

Self vs. non-self discrimination is a fundamental process across biological systems, from immune recognition in multicellular organisms to CRISPR-based defense in bacteria^1^. Similarly, social animals have evolved mechanisms to recognize and discriminate against individuals outside of their group or colony to defend themselves against exploitation. In social insects, this discrimination relies on complex mixtures of cuticular hydrocarbons (CHCs), chemical compounds with low volatility that individuals carry on the body surface^2–6^. The role of hydrocarbons in nestmate recognition and non-nestmate discrimination has been documented across various social insect species, e.g., via experiments measuring aggression toward nestmates artificially treated with synthetic hydrocarbons^6–9^ or toward neutral objects coated with either natural hydrocarbon extracts or synthetic blends mimicking colony odors^10–13^. Discrimination can also arise when key components of the CHC profile are missing^14^.

The CHCs of social insects belong to different structural classes, such as linear alkanes, methyl-branched alkanes, and alkenes^5,8,15^. Different species often differ in their CHC profiles and the types of compounds they produce. Variation in CHC profiles between workers from different colonies is also ubiquitous within species, although here it usually involves differences in the relative abundance of shared hydrocarbons^5,16^. While CHC profiles are largely genetically determined^17^, a uniform odor emerges for each colony because nestmates exchange CHCs via mechanisms like trophallaxis (exchange of regurgitated liquid food), allogrooming (cleaning of one individual by another with legs and mouthparts), and contact with a shared nest substrate^3,17–20^. Newly emerged (callow) social insect workers initially only carry small amounts of CHCs, and then synthesize their own CHCs but also acquire hydrocarbons from nestmates, gradually acquiring the colony odor^17,21–24^. Young workers learn to recognize the colony odor during the first few days of adult life^17,25^, and because the colony odor can change over time due to changes in diet or nesting substrate, individuals should be able to continuously update their internal representation of the colony odor^14,17,25–29^. Supporting the idea that early social exposure shapes discrimination abilities, studies have shown that workers reared in experimental mixed-species groups exhibit reduced interspecies aggression compared to individuals who have never encountered the other species. This shows that ants can learn and recognize a wide range of colony odors, including both conspecific and heterospecific ones^25,30–32^. However, we know little about how flexible CHC recognition cues and the learned odor templates are later in life, and over what timescales changes can occur. Addressing these questions has been challenging because it requires precise experimental control over an individual’s genotype and its social environment, as well as the ability to identify and follow the same individual across different social contexts.

Here, we use the clonal raider ant *Ooceraea biroi* to study changes of CHC profiles and plasticity of nestmate recognition “templates”. *O. biroi* colonies lack queens and consist entirely of totipotent workers that reproduce asexually^33–35^. All workers in natural colonies are clonally identical, and colonies from one clonal genotype recognize and attack ants from other clonal genotypes^36^. Furthermore, ants from different clonal lines can be mixed in experimental colonies. Together, this offers exceptional control over the genotype of individuals and the genotypic composition of social groups. In addition, molecular genetic tools have been established in this species, including transgenic lines that express fluorescent markers. This allows us to track individuals across different social environments. Using a cross-fostering design, we tested whether ants can acquire new CHC profiles from foster colonies and studied the temporal dynamics and plasticity of the ants’ behavioral responses to nestmates and non-nestmates of different genotypes.

## RESULTS

### Aggression toward foreign genotypes is stable over time

For this study, we used a strain derived from wildtype clonal line B that carries a transgenic insertion of the red fluorescent protein dsRed (referred to as Bˈ)^35^. These ants show strong red fluorescence broadly across their body, making them easily identifiable under a fluorescence microscope (**Figure S1A**). We first examined the baseline levels of aggression that these ants display toward individuals from different clonal genotypes (**Figure 1A**). We established host colonies of 25 Bˈ ants and 13 larvae and introduced one focal ant from the same or a different clonal genotype at a time (genotype Bˈ, as well as wild-type genotypes A, B, and D) (**Figure 1B**). Bˈ ants were more aggressive toward focal ants from genotypes A and D than toward Bˈ and B ants, demonstrating that they can effectively discriminate between nestmates and non-nestmates of different clonal genotypes (**Figure 1C**). The lack of aggression of Bˈ toward line B generalizes across different line B stock colonies (**Figure S1B**) and is expected because the dsRed transgenic line was created in a wildtype line B genetic background^35^. Host aggression toward focal ants was not measurably affected by reciprocal aggression, as the number of aggressive behaviors by hosts did not correlate with the number of aggressive behaviors performed by focal ants (**Figure S1C**).

**Figure 1:**
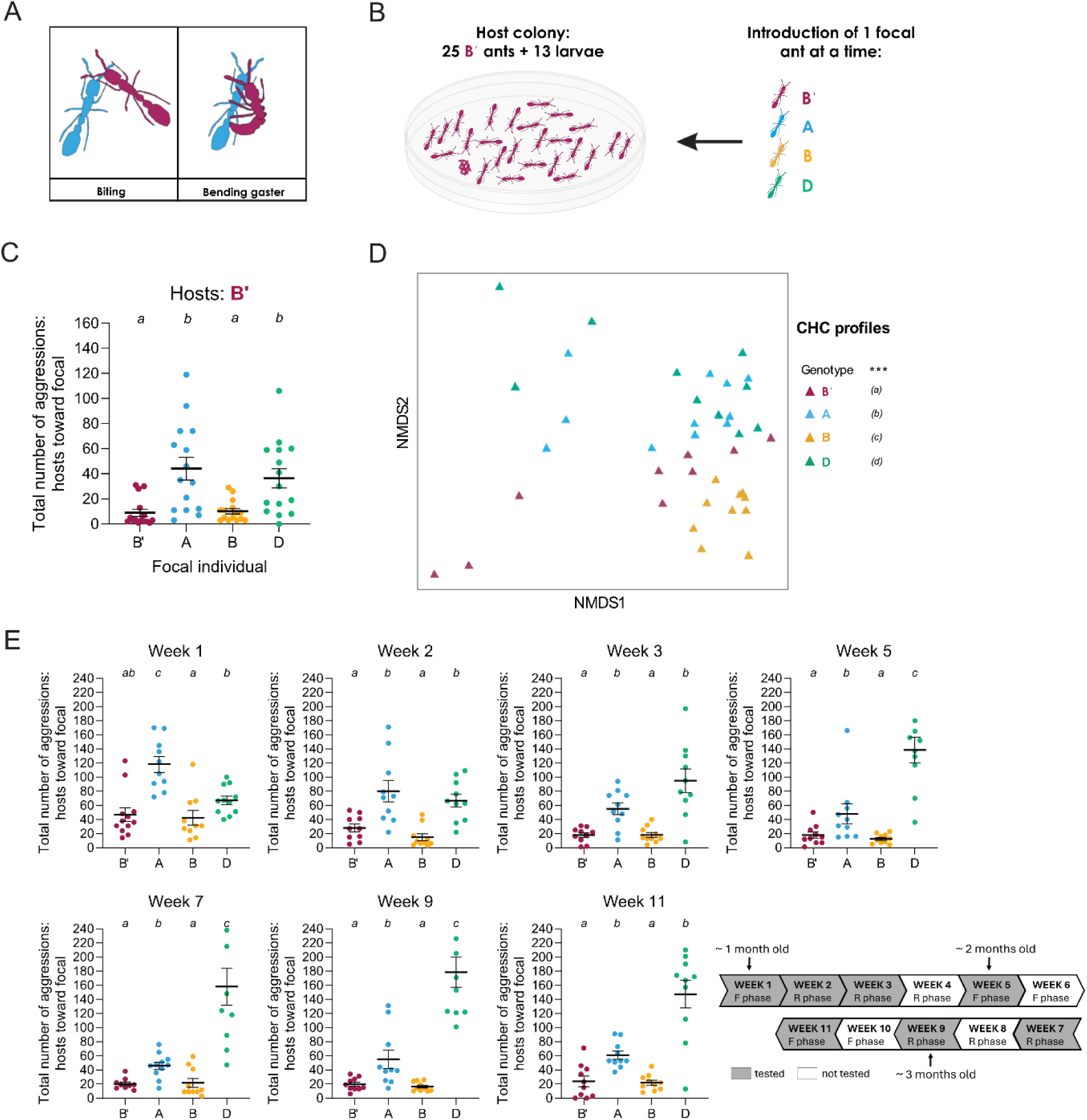
Aggression toward foreign genotypes is stable over time. (A) Types of aggressive behaviors measured in the clonal raider ant. Biting: a host ant bites the focal individual’s head, thorax, abdomen, or appendages (antennae or legs). This can result in dragging the focal ant around the arena by an antenna or leg. Bending gaster: the host ant curls its gaster toward the focal individual. While the significance of this behavior is not clear (it might represent an attempt to sting), it is a strong indicator of aggression. The different aggressive behaviors are shown in **Video S1**. (B) Schematic of the non-nestmate discrimination assay. Focal ants of genotypes Bˈ, A, B and D were introduced one at a time to a host colony composed of 25 Bˈ ants and 13 larvae. The aggressive behaviors exhibited by host ants toward the focal individual during 30 minutes were counted. (C) Total number of aggressive behaviors exhibited by Bˈ host ants toward focal ants of different genotypes. N=15 replicates per genotype. Error bars: standard error of the mean (SEM). Different letters indicate statistically significant differences between focal genotypes (p < 0.05). (D) NMDS analysis of CHC profiles from ants of genotypes Bˈ (magenta), A (blue), B (yellow) and D (green). Data points represent individual ants (N=9-11 replicates per genotype). The NMDS analysis was conducted based on the relative abundance of each CHC compound across the different genotypes (a list of the 24 *O. biroi* compounds, along with their relative abundances, is provided in **Supplementary Table S1**). *** indicates an overall significant difference among genotypes (p < 0.001), and different letters indicate CHC profile differences (multiple comparisons) between genotypes. (E) Total number of aggressive behaviors exhibited by original (not cross-fostered) Bˈ ants toward focal ants of genotypes Bˈ, A, B and D over time. These behavioral assays were performed to measure variability of aggressive behaviors over time and control for potential age-dependent (1-vs. 2- vs. 3-month-old ants) and cycle phase-dependent (F: foraging phase; R: reproductive phase) effects, as illustrated in the timeline cartoon. N=9-12 replicates per genotype. Error bars: standard error of the mean (SEM). Different letters indicate differences between focal genotypes (p < 0.05). Details of statistical analyses are provided in **Supplementary Table S2.**

To measure chemical differences between different genotypes and between different clonal line B stock colonies, we extracted CHCs from ants and analyzed them using gas chromatography-mass spectrometry (GC-MS). Similar to previous work^37^, ants from different clonal genotypes differed in the relative abundance of the 24 CHC compounds found in *O. biroi* (**Figure 1D; Figure S1D-F**). This is consistent with the ants’ ability to distinguish between different genotypes. Despite detectable differences in the CHC profiles of Bˈ colonies and some of the different clonal line B stock colonies (**Figure 1D; Figure S1G**), ants from these various colonies did not seem to recognize each other as non-nestmates in our behavioral assays (**Figure 1C; Figure S1B**).

Colonies of clonal raider ants undergo stereotypical behavioral and reproductive cycles, alternating between ca. three week-long ‘reproductive phases’ in which the ants lay eggs and ca. two week-long ‘foraging phases’ in which colonies contain larvae that the ants provision with food. We therefore conducted an experiment to assess the variability of aggression levels over time, accounting for the ants’ age and physiological state across different phases of the colony cycle. For this experiment, we set up a host colony of one month old Bˈ ants and measured their aggression toward ants of different genotypes over several weeks (experiments conducted in weeks 1, 2, 3, 5, 7, 9 and 11). We conducted behavioral assays on two consecutive days each week, with a five-day recovery interval between tests in consecutive test weeks. Regardless of age and cycle phase, the ants were consistently aggressive toward focal individuals of genotypes A and D (**Figure 1E**). This shows that, under this exposure regimen, the ants’ baseline discriminatory behavior is stable across time (at least from ca. 1 month to 3.5 months of age) and not dependent on the behavioral and physiological stage of the colony.

### Prolonged exposure establishes tolerance toward foreign genotypes

To explore the plasticity of non-nestmate discrimination, we investigated whether Bˈ ants housed in a foster colony of a different clonal genotype develop a modified CHC profile, and whether prolonged exposure to foster ants alters their aggressive behavior toward individuals of the foster colony’s genotype. We cross-fostered Bˈ callows (2 to 4 days old) for one month in colonies from three different clonal genotypes: one that does not elicit aggression from Bˈ ants (genotype B) and two that do elicit aggression from Bˈ ants (genotypes A and D). After one month, Bˈ ants were separated from their respective foster colony and combined in new colonies composed entirely of Bˈ ants that had spent their early adulthood either in line B colonies, line A colonies, or line D colonies (**Figure 2A**).

**Figure 2:**
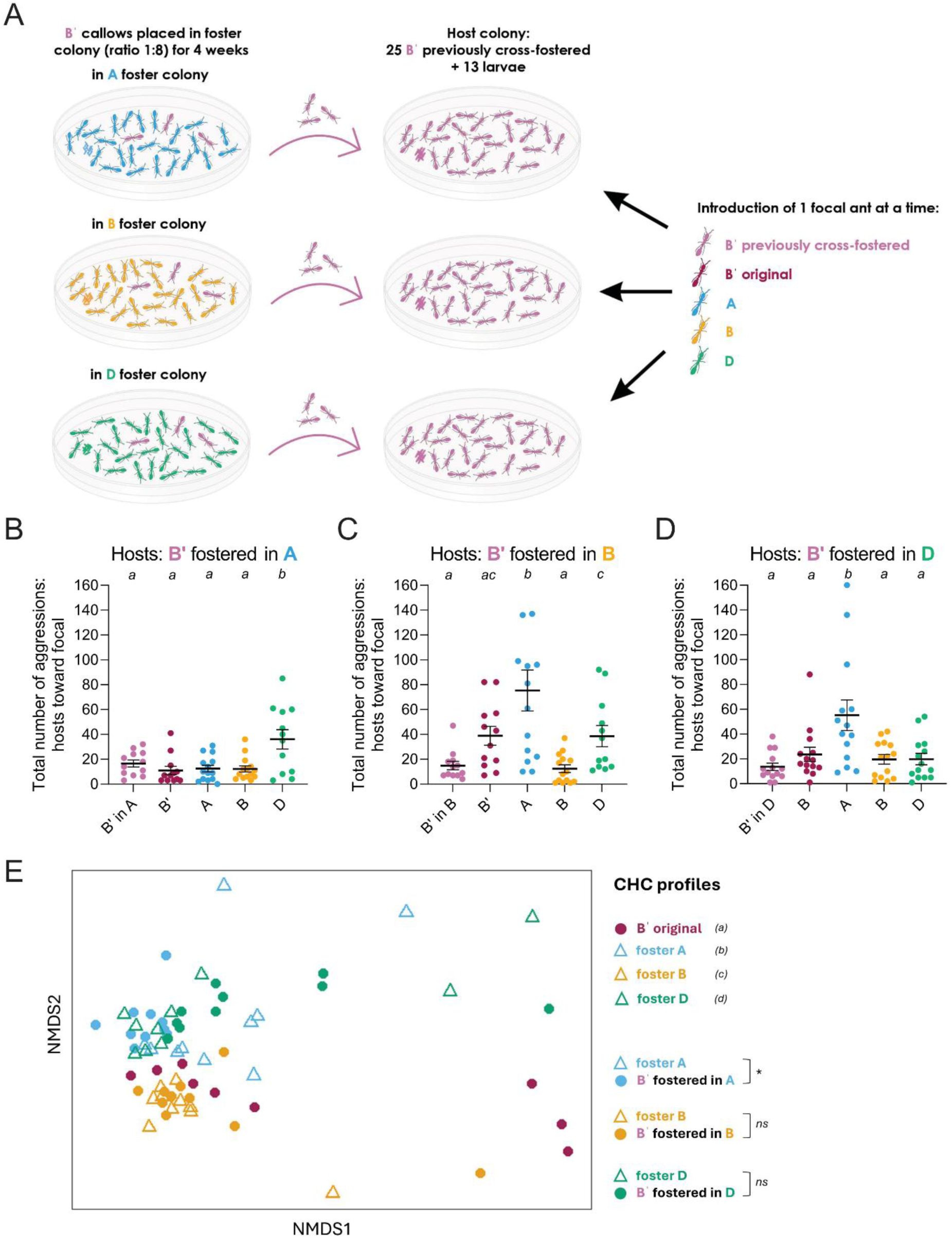
Prolonged exposure establishes tolerance toward foreign genotypes. (A) Schematic of the behavioral assay. Bˈ callows were placed and kept in foster colonies from different genotypes (i.e. A, B and D) for 1 month in a ratio of 1 Bˈ ant to 8 foster ants. Bˈ ants were then separated from foster colonies and grouped to make host colonies of 25 Bˈ ants and 13 larvae. Focal ants of original Bˈ, Bˈ raised in a foster colony, as well as A, B and D wild-type genotypes were introduced one by one in Bˈ host colonies, and aggressive behaviors exhibited by host ants toward respective focal ants during 30 minutes were counted. (B)(C)(D) Total number of aggressive behaviors exhibited by Bˈ ants raised in foster colonies of genotypes B, A and D, respectively, toward focal ants of each genotype. N=12-14 replicates per genotype. Error bars: standard error of the mean (SEM). Different letters indicate differences between focal genotypes (p < 0.05). (E) CHC profiles of foster wild-type ants (empty triangles) and Bˈ ants (filled circles) visualized with an NMDS analysis. The original Bˈ ants are colored magenta, while Bˈ raised in A, B and D are in blue, yellow and green, respectively. Data points represent individual ants (N=9-10 replicates per genotype). The NMDS analysis was conducted based on the relative abundance of each CHC compound across the different groups. Different letters indicate significant differences in CHC profiles between genotypes (multiple comparisons). Statistical comparisons between foster wild-type ants and cross-fostered Bˈ ants are also indicated (ns: not significant, *: p < 0.05). Details of statistical analyses are provided in **Supplementary Table S2.**

Bˈ ants fostered in B colonies were not more aggressive toward original (not cross-fostered) Bˈ ants or B ants than they were toward other Bˈ ants fostered in B colonies (**Figure 2C**). However, as expected, they behaved aggressively toward A and D ants (**Figure 2C**). Importantly, Bˈ ants fostered in A and D colonies showed no elevated aggression toward focal ants of their respective foster genotype or of their own genotype but maintained aggression toward ants from the opposite genotype (D and A, respectively) (**Figure 2B&D**). The genotype of the foster colony thus altered the pattern of aggression displayed by Bˈ ants, revealing that prolonged exposure to ants of a different genotype promotes tolerance.

Following behavioral experiments, we analyzed the chemical profiles of experimental ants using GC-MS and found that the CHC profiles of Bˈ ants had shifted in response to the foster colony environment. Specifically, the CHC profiles of these ants differed from those of the original Bˈ ants and had become more similar to the foster colony’s CHC profile (**Figure 2E**). Our results suggest that exposing Bˈ ants to foster ants for one month is sufficient for them to acquire a modified CHC profile and to learn the gestalt odor of the foster colony, demonstrating that both CHC profiles and discrimination behavior change in adult ants.

Bˈ ants fostered in A, B and D colonies did not show increased aggression toward the original Bˈ ants (**Figure 2B-D**). This suggests that, although cross-fostered Bˈ ants are consistently exposed to a different CHC profile and even acquire a different CHC profile themselves, this does not result in aggression toward their genetically encoded “default” CHC profile. To determine whether this lack of discrimination is due to the ants’ experience in Bˈ colonies at the larval, pupal, or very early adult stage, we cross-fostered Bˈ eggs in an A colony and waited until the resulting ants were adults and had reached one month of age. At that point, the Bˈ ants were separated from the foster ants, grouped together to form a host colony, and tested for aggressive behavior toward focal ants of different genotypes (**Figure S2A**). While aggression toward D ants was elevated as expected, the level of aggression toward all other focal genotypes was low and not statistically different from the control (**Figure S2B**). In other words, although the CHC profile of Bˈ ants raised in the A foster colony differed from that of the original Bˈ ants (**Figure S2C**), these ants were not aggressive toward the original Bˈ focal ants, despite never having encountered them before (**Figure S2B**). This indicates that ants do not learn a colony’s odor during the larval or pupal stage but instead suggests that some other mechanism prevents them from becoming aggressive toward their genetically encoded CHC profile. For example, it might be possible that ants smell and adapt to their intrinsic CHC profile even in situations where this profile differs from the odor of their colony. We also cross-fostered Bˈ eggs in a B foster colony (**Figure S3D**) and found that, although Bˈ ants raised in the B colony had no prior contact with original Bˈ ants, they showed no aggression toward either B or original Bˈ individuals but were aggressive toward A and D focal ants (**Figure S2E**). Here again, aggression patterns were similar to those observed when Bˈ ants were cross-fostered as adults, confirming that the lack of aggression toward Bˈ and B in our cross-fostering experiments does not stem from exposure during the larval or pupal stage, or the first hours of adult life.

### Tolerance degrades with prolonged separation

Having established that Bˈ ants fostered in A colonies acquire a CHC profile similar to A and initially exhibit reduced aggression toward A ants after having been separated from their foster colony (**Figure 2C**), we asked whether this change persists following separation. For that, we cross-fostered Bˈ callows in line A colonies until they were one month old. We then separated them from the line A foster colonies and pooled them into new colonies composed exclusively of Bˈ ants that had previously been fostered with line A. We measured their CHC profiles and aggression toward ants of various genotypes, including ants from the line A foster colony, using a paradigm directly comparable to the experiment reported in **Figure 1E** (i.e., at 1, 2, 3, 5, 7, 9 and 11 weeks after separation; **Figure 3A**).

**Figure 3:**
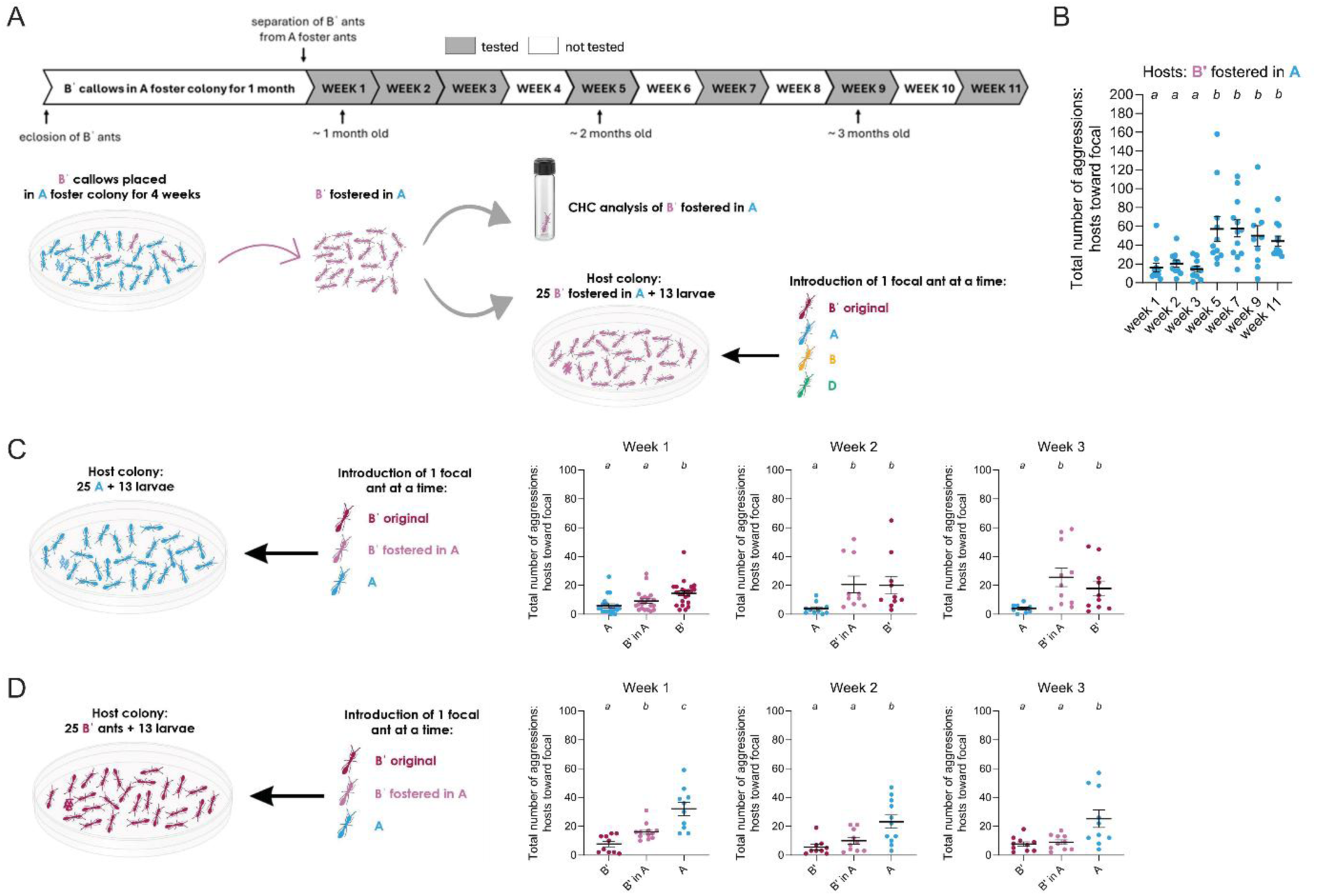
Tolerance degrades with prolonged separation. (A) Schematic of the behavioral and chemical assays. Bˈ callows were placed and kept in an A foster colony for 1 month in a ratio of 1 Bˈ ant to 8 A foster ants. Bˈ ants were then separated from foster colonies. Some were grouped to form host colonies of 25 Bˈ ants and 13 larvae for behavioral experiments, while the rest were kept in a separate dish for chemical analyses. Focal ants of genotypes Bˈ, A, B and D were introduced one at a time to Bˈ host colonies and the aggressive behaviors exhibited by host ants toward the respective focal ant during 30 minutes were counted. Behavioral assays (see Figure 3B) were conducted in weeks 1, 2, 3, 5, 7, 9 and 11 after separation from the foster colony, and chemical assays (see **Supplementary Figure S3D**) were conducted in weeks 1, 2, 3 and 5 after separation from the foster colony. (B) Total number of aggressive behaviors exhibited by Bˈ ants raised in A foster colonies toward focal ants of genotype A over time. Other focal genotypes are shown in **Supplementary Figure S3A**. N=10-12 replicates per week. Error bars: standard error of the mean (SEM). Different letters indicate differences between weeks (p < 0.05). (C) Original Bˈ ants, Bˈ ants raised in A colonies, and A ants were introduced one at a time into an A host colony of 25 ants and 13 larvae, and the aggressive behaviors exhibited by the host ants toward the respective focal ant during 30 minutes were counted. These behavioral assays were conducted at various time points, alongside assays using Bˈ ants raised in A colonies as host ants (see Figure 3A**&B**). Results from behavioral experiments during weeks 1, 2 and 3 after B’ ants had been separated from their A foster colony are shown here, and results from weeks 5, 7 and 9 are shown in **Supplementary Figure S3B**. N=10-23 replicates per genotype. Error bars: standard error of the mean (SEM). Different letters indicate differences between focal genotypes (p < 0.05). (D) Original Bˈ ants, Bˈ ants raised in A colonies, and A ants were introduced one at a time into a Bˈ host colony of 25 ants and 13 larvae, and the aggressive behaviors exhibited by the host ants toward the respective focal ant during 30 minutes were counted. These behavioral assays were conducted at various time points, alongside assays using Bˈ ants raised in A colonies as host ants (see Figure 3A**&B**). Results from behavioral experiments during weeks 1, 2 and 3 after B’ ants had been separated from their A foster colony are shown here, and results from weeks 5, 7 and 9 are shown in **Supplementary Figure S3C**. N=9-12 replicates per genotype. Error bars: standard error of the mean (SEM). Different letters indicate differences between focal genotypes (p < 0.05). Details of statistical analyses are provided in **Supplementary Table S2.**

Cross-fostered Bˈ ants were not aggressive toward A ants during weeks 1, 2 and 3, but they responded aggressively from week 5 onward (**Figure 3B**). This indicates that cross-fostered ants continued to treat line A ants as nestmates for at least three weeks before reverting to recognizing them as non-nestmates by week 5. These results imply that tolerance toward the foster genotype declined after several weeks of separation from the foster colony. Additional aggression assays with other genotypes revealed low aggression toward Bˈ and B ants and high aggression toward D ants across all time points (**Figure S3A**), indicating that the loss of non-nestmate discrimination in B’ ants was specifically toward genotype A ants.

We then asked how CHC profiles of formerly cross-fostered ants change once they have been separated, and how this might affect the aggression they receive from ants of their former host genotype as well as ants of their own genotype. After one week of separation, host ants from genotype A were not aggressive toward cross-fostered Bˈ ants, while they were aggressive toward original Bˈ ants, as expected (**Figure 3C**). However, from week 2 onward, A hosts displayed aggression toward Bˈ ants fostered in A at levels similar to those shown toward original Bˈ ants (**Figure 3C, Figure S3B**). This suggests that the changes in CHC profiles of cross-fostered ants within the first two weeks after separation were sufficient for their former hosts to recognize them as non-nestmates. Consistent with this timeline, Bˈ host ants were aggressive toward Bˈ ants that had been cross fostered in line A colonies one week after separation (**Figure 3D**). However, from two weeks onward, aggression toward formerly cross-fostered ants had subsumed (**Figure 3D, Figure S3C**). This suggests that cross-fostered Bˈ ants had restored the decisive features of their original CHC profile within two weeks of separation from their line A foster colonies.

Chemical analyses did not precisely mirror the behavioral dynamics. While the CHC profiles of Bˈ ants had already shifted one week after separation from the foster colony and kept changing throughout the experiment, they never fully reverted to the original Bˈ profile even five weeks after separation (**Figure S3D**). This illustrates that statistical analyses of entire CHC profiles are not perfectly predictive of the ants’ behavioral responses, possibly because behavioral discrimination relies on the ratio of specific subsets of CHC cues rather than on the entire CHC profile. In other words, our chemical analyses might also detect differences in CHC profiles that the ants either cannot sense or that are not relevant for nestmate discrimination.

### Low-level sporadic exposure is sufficient to maintain tolerance

The experiments described in **Figure 3A** were conducted weekly during the first three weeks, and then biweekly until week 11. Bˈ ants that had initially been fostered with line A first exhibited aggression toward genotype A at 5 weeks after separation. However, because this timepoint followed the first two-week interval without exposure, the reappearance of aggression could be due to the total time elapsed since separation or the increased latency between re-exposure events. To distinguish between these possibilities and to test whether repeated exposure is required to maintain the lack of aggression toward the foster genotype, we conducted an additional experiment spanning the first seven weeks after separation from the foster colony. As before, Bˈ callows were cross fostered in an A colony for 4 weeks. After separation, they were pooled in a group of 25 to form a host colony, which was then tested for aggression. Tests were conducted every week for the first five weeks. As in previous experiments, behavioral assays were conducted on two consecutive days each week, with a five-day break in between. To test the effect of interrupted exposure, the host colony was then not tested in week 6 and retested in week 7 (**Figure 4A**).

**Figure 4:**
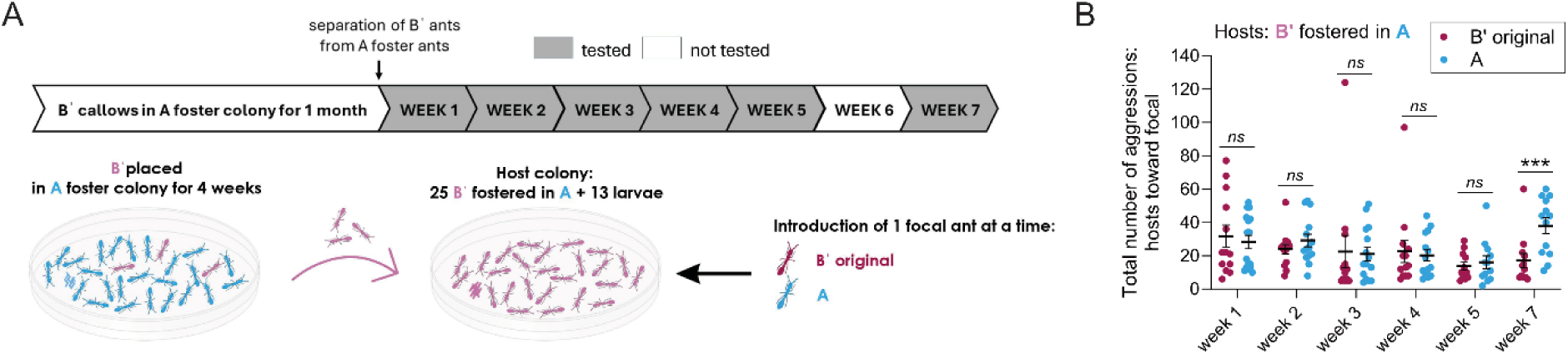
Low-level sporadic exposure is sufficient to maintain tolerance. (A) Schematic of the behavioral assay. Bˈ callows were kept in genotype A foster colonies for 1 month in a ratio of 1 Bˈ ant to 8 A ants. Bˈ ants were then separated from their foster colonies and combined into a host colony of 25 Bˈ ants and 13 larvae. Original Bˈ ants and A ants were introduced one at a time into this Bˈ host colony, and the aggressive behaviors exhibited by the host ants toward the respective focal ant during 30 minutes were counted. Behavioral assays were conducted in weeks 1, 2, 3, 4, 5 and 7 after Bˈ ants had been separated from their genotype A foster colonies. (B) Total number of aggressive behaviors exhibited by Bˈ ants raised in genotype A foster colonies toward focal ants of genotypes Bˈ and A. N=11-16 replicates per genotype. Error bars: standard error of the mean (SEM). ns: not significant, ***: p < 0.001. Details of statistical analyses are provided in **Supplementary Table S2.**

Bˈ ants that had initially been fostered in line A colonies showed no elevated aggression toward A ants relative to other Bˈ ants in weeks 1, 2, 3, 4 and 5 after separation (**Figure 4B**). However, they were aggressive toward A ants in week 7, suggesting that skipping exposure to A ants during week 6 was sufficient to restore non-nestmate discrimination (**Figure 4B**). This shows that repeated exposure is required to maintain amicable interactions and to prevent aggression between cross-fostered ants and their former hosts. Exposure during behavioral experiments was limited to two days per week and interrupted by five days of recovery without exposure. This indicates that, once behavioral tolerance has been established, it can be sustained by even sporadic contact with foreign genotypes.

## DISCUSSION

### Plasticity in behavioral responses to colony odors

Bˈ ants housed in foster colonies of different genotypes from a young age showed reduced aggression toward individuals of those genotypes. Similar effects of early social experience on aggression have been observed in ants reared in mixed-species groups, which display reduced aggression toward heterospecifics^25,30–32^. Our results confirm that the neural colony odor “template” is plastic in young ants and that it is shaped by the social environment.

Following separation from the foster colony, Bˈ ants became aggressive toward the foster genotype within weeks unless they were sporadically re-exposed. This shows that odor-based discrimination is not fixed in older ants either, and that it depends on ongoing social experience and sensory reinforcement. These findings highlight the importance of exposure in maintaining low aggression and social tolerance once it has been established. Interestingly, a recent study on black garden ants (*Lasius niger*) reported that ants attacked by non-nestmates learn the chemical cues of their aggressors and become more hostile in later encounters^38^. Thus, while repeated peaceful exposure reduces aggression in our study, hostile interactions reinforce discrimination and increase defensive responses. This contrast underscores that the direction and persistence of behavioral plasticity in non-nestmate discrimination may critically depend not only on encounter frequency, but also on whether the experienced encounters are amicable or antagonistic.

Our experiments revealed an additional novel facet of non-nestmate discrimination. Bˈ ants raised in foster colonies from the egg stage were continuously exposed to the A genotype and had no prior contact with pure Bˈ individuals. Yet, despite having never encountered ants of their own original genotype, they did not show aggression toward them. This suggests that some mechanism prevents aggression toward the ants’ genetically determined recognition cues. For instance, ants may sense and learn or become accustomed to their own intrinsic hydrocarbon profile, in addition to the colony’s odor.

This dual influence of genetics and experience implies a distinction between colony- and individual-level recognition. Previous studies have shown that ants can become aggressive toward members of their own colony when dietary or other environmental differences result in altered CHC profiles^14,17,25–29,39^. In our experiments, Bˈ ants were maintained in a foster environment under standardized dietary and environmental conditions, and their CHC profiles mostly, if not exclusively, changed through contact with foster nestmates^3,21^. While this is an example of the (social) environment shaping CHC profiles, unlike dietary shifts, cross-fostering unlikely changed the CHC composition that Bˈ ants themselves synthesize. As a result, these ants might have remained exposed to the chemical bouquet of their original genotype by detecting their own endogenous odor. Consequently, in the experiments reported here, the tolerance of cross-fostered ants toward CHC cues appears to broaden rather than shift.

### The neural basis of non-nestmate discrimination

Behavioral assays were conducted on two consecutive days each week, with a five-day break in between (see STAR Methods). Surprisingly, this sporadic exposure to the foster genotype was sufficient to maintain behavioral tolerance. This suggests that ants have some sort of “memory” of their former colony’s odor, raising questions about the underlying neural mechanisms.

Historically, non-nestmate discrimination has been thought to rely on learning the CHC profile of nestmates during early social interactions, creating a neural template stored in brain regions such as the mushroom bodies^2,3,9,31,40,41^. Such a memory-based system would have to be flexible, allowing templates to be updated throughout an ant’s life in response to changes in colony odor and social context, such as repeated encounters with friends or foes^5,15,32,38,42^. Discrimination between nestmates and non-nestmates occurs when an ant compares the detected chemical label of another individual to its internal template. If the label matches the template that individual is treated amicably, while a mismatch triggers aggression^2,3,8,9,25,40^. The decision to behave aggressively is often modeled as a threshold process, where aggression is triggered if the perceived chemical difference exceeds a certain threshold, while individuals below that threshold are accepted or ignored^2,17,24^.

In contrast, sensory adaptation has been suggested as a simpler alternative mechanism that does not require canonical learning or memory^9^. When ants are repeatedly exposed to a non-threatening odor, their sensory system becomes less responsive to that stimulus. As a result, rather than forming an active memory, the ant’s sensory neurons or downstream neural circuits become less reactive, leading to decreased aggression^2,9,14,43–46^. For example, sensory adaptation at the periphery, such as desensitization of olfactory receptor neurons in the antennae, has been implicated in the process^41,47^. Such peripheral filtering could allow ants to ignore familiar colony odors while remaining sensitive to unfamiliar and foreign profiles indicative of non-nestmates. In this model, discrimination occurs even in the absence of an explicit central template: ants simply fail to detect nestmates as distinct stimuli but respond to non-nestmates.

Consistent with this possibility, empirical studies suggest that ants accept individuals by default if they lack undesirable cues. In other words, ants recognize foes rather than friends, and aggression seems to be specifically elicited by unfamiliar or ’foreign’ odors, rather than by the absence of familiar ones^9,39,47^. This implies that ants primarily perform non-nestmate discrimination, rather than nestmate recognition, a process that might not strictly require a complex template stored in long-term memory.

Our study highlights the importance of repeated exposure in maintaining tolerance of former nestmates. However, these exposures can be sporadic, with up to five intermittent days without exposure. Sensory adaptation usually operates over shorter timescales and decays rapidly once stimulation stops. For example, olfactory adaptation generally returns to baseline within minutes after stimulus removal in *Drosophila* ^48,49^, and within hours in *Caenorhabditis elegans*^50^. Memory-based processes, on the other hand, can sustain behavioral changes over longer periods, and ants are known to form olfactory memories lasting from hours to months^31,51–53^. For example, *Formica fusca* workers retain odor-reward associations for up to three days^51^, *Cataglyphis fortis* can remember food odors for nearly a month^52^, and *Pachycondyla villosa* and *P. inversa* queens recall individual identities based on odor after 24 hours of separation^53^. Long-term memory has also been implicated in ant nestmate recognition, as workers reared in mixed-species groups tolerate familiar individuals from a different species even after one year of separation^31^. These examples underscore that olfactory memories in ants can be remarkably persistent. Based on these considerations, sensory adaptation alone seems unlikely to explain the persistence of tolerance to different clonal genotypes over several days without exposure in the clonal raider ant, suggesting that longer-lasting memories are also involved.

## Conclusion

Our experiments demonstrate that non-nestmate discrimination in ants is highly plastic and responsive to the social environment. While ants require prolonged exposure to develop tolerance toward foreign genotypes, once established, this tolerance can be maintained via sporadic encounters. At the same time, aggression returns to baseline within two weeks without repeated exposure. This plasticity in non-nestmate discrimination implies that some form of underlying learning and memory is active throughout adult life. However, the neural underpinnings of these phenomena remain unknown. This study, in concert with recently established neurogenetic tools for the clonal raider ant^60–62^, opens the door to investigate the neurobiological basis of non-nestmate discrimination and the associated behavioral plasticity in social insects at the circuit level.

The dynamic nature of tolerance we observe in ants shows intriguing parallels to how other biological and social systems manage discrimination of “non-self” from “self”. In the vertebrate immune system, for example, encounters with foreign antigens can lead either to immune activation or to tolerance, depending on the timing, dose, and context of exposure^54,55^. Repeated low-level exposure, such as during allergen immunotherapy, promotes long-lasting tolerance through active regulatory mechanisms^56,57^. Similarly, social psychology studies indicate that early and repeated exposure to members of other human groups can reduce xenophobic attitudes later in life^58,59^, supporting the idea that continuous, non-threatening social contact promotes tolerance toward “foreigners”. Across these diverse systems, tolerance emerges from prolonged and repeated exposure, and requires recognition, memory, and reinforcement.

## MATERIALS AND METHODS

### Ant husbandry and maintenance

For this study, we used *Ooceraea biroi* stock colonies of clonal lines A, B and D that were originally collected in Okinawa (Japan), St. Croix (U.S. Virgin Islands) and Bangladesh, respectively^63,64^. Prior to this study, all stock colonies had been maintained in the laboratory under standardized conditions for at least ten years. Additionally, we used a line derived from clonal line B that carries a transgenic insertion of the red fluorescent protein dsRedB^60^. We refer to this line as Bˈ here. Bˈ ants can be easily distinguished and separated from wild-type ants under an epifluorescence microscope (**Figure S1A**). Colonies were maintained in environmental rooms at 25°C and 60% humidity in Tupperware containers with a plaster of Paris floor. The plaster was kept humid and colonies were fed with frozen fire ant pupae three times per week during the foraging phase.

### Cross-fostering protocol

#### Fostering Bˈ callows

Callows (newly eclosed ants, 2 to 4 days old) were used for cross-fostering because, as in other ant species, they can be transferred across colonies without eliciting aggression^31^. Callows have only small amounts of hydrocarbons on the cuticle (**Figure S4A**). Chemical analyses further revealed that the CHC profiles of 1-day- and 1-week-old callows significantly differed from that of 1- and 3-month-old adults (**Figure S4B**), which likely explains the tolerance by resident ants.

Bˈ callows were placed in genotype A, B or D foster colonies in a ratio of 1 Bˈ callow to 8 adult foster ants of unknown age for 4 weeks. Foster colonies were maintained as described above. After four weeks, the colonies were examined under an epifluorescence microscope and Bˈ ants, now four weeks old, were separated from the foster ants. Bˈ ants were grouped together into a new colony, which was then used for experiments as described in the ‘Behavioral assays’ section.

#### Fostering Bˈ eggs

Bˈ eggs were placed in genotype A and B foster colonies in a ratio of 1 Bˈ egg to 8 adult foster ants of unknown age. Eggs from wild-type foster ants were simultaneously added to maintain a natural ratio of eggs to adults, which increases colony stability and egg survival. Once adults from this cohort of eggs had eclosed, some of the older foster ants were removed to maintain a ratio of 1 Bˈ ant to 8 foster ants. Four weeks later, i.e., once the B’ ants were four weeks old, Bˈ ants were separated from the foster ants under an epifluorescence microscope and grouped into a new colony that was then used for experiments as described in the ‘Behavioral assays’ section.

### Behavioral assays

#### Preparation of host colonies and focal individuals

Three days before behavioral experiments, we set up experimental ‘host’ colonies composed of 25 ants and 13 larvae in standard Petri dishes (50 x 9mm) with a moist plaster of Paris floor. In each case, adult ants and larvae were derived from the same laboratory stock colony or foster colony during the foraging phase of the colony cycle. Each of the 25 host ants was painted with a color dot (Sanford® Uni® Paint oil-based markers) on the abdomen. Focal individuals were also collected from stock colonies and foster colonies during the foraging phase and were marked with a different color before being transferred to separate Petri dishes with moist plaster of Paris floors. To exclude any possible bias related to the paint colors assigned to host and focal individuals from different colonies, we randomized the color assignments across experiments. Petri dishes containing host colonies and focal individuals were then stored at 25 °C with food for two to three days until the experiments began.

For some genotypes, we used focal ants from different stock colonies for different experiments. To ensure that this did not influence aggressive behavior and non-nestmate discrimination, we grouped 25 Bˈ ants with 13 larvae into a host colony and measured aggression toward focal ants from 4 different B colonies (B1-B4) (**Figure S1B**). Despite variation in CHC profiles among different B colonies (**Figure S1G**), Bˈ ants displayed no aggression toward any B focal ants, while being aggressive toward genotype A ants (**Figure S1B**). This indicates that using ants from different stock colonies of the same clonal genotype does not measurably affect behavioral outcomes in our experiments.

#### Experimental arenas and procedures

All behavioral experiments were conducted directly in the host colony’s Petri dish. This minimizes perturbation of the host colony and preserves the local chemical environment. Focal individuals were introduced one at a time. Immediately after introducing the focal ant, the Petri dishes were placed in a ventilated custom enclosure from the company Loopbio (46(D) x 46(L) x 52(H) cm) with built-in temperature control (set to 25°C) and LED lighting. Videos were recorded with Basler a2A4504-18ucPRO cameras for 30 minutes. After each 30 minute assay, the focal individual was removed and, after a 5-minute break, a new focal individual from a different genotype was added. Two to three host colonies were used in each experiment, i.e., each host colony was tested repeatedly with different focal individuals. Each focal individual, however, was used only once to avoid potential transfer of host CHCs to focal ants, which could modify the hosts’ behavior in subsequent assays. Forceps were washed with ethanol between assays to avoid transfer of CHCs or other odorants. To prevent an effect of day and time of day, assays were spread out over multiple days. Aggression was measured by counting instances of aggressive behavior of host ants toward the focal individual (details below).

For the behavioral experiments spanning multiple weeks, assays were performed on two consecutive days each week, with a five-day break in between. Host colonies were maintained in the same dishes with food under controlled environmental conditions between testing sessions.

#### Non-nestmate discrimination

Non-nestmate discrimination assays were performed by manually counting the total number of aggressive behaviors exhibited by host ants toward the focal individual during 30 minutes. We counted instances in which a host ant bit the focal ant’s body or appendages (legs and antennae), as well as instances in which a host ant bent her abdomen toward the focal ant (**Figure 1A; Video S1**). All behavioral observations were conducted blindly with respect to the focal ant’s genotype or colony of origin. Bˈ transgenic ants are not less aggressive toward ants of different genotypes than wild-type B ants (**Figure S4C**), showing that their ability to discriminate against non-nestmates has not been altered.

### Chemical analysis

Ants for chemical analyses were frozen at -20°C, and CHCs were extracted from single frozen ants by immersing them in 200µL hexane for 10 minutes. Between 8 and 10 individual ants (i.e., 8-10 replicates) were extracted per genotype/condition. Each ant was in the foraging phase when frozen, eliminating potential confounding effects of variation in CHC profiles throughout the colony cycle. CHC extracts were then evaporated to a volume of approximately 15µL, of which 1µL was analyzed using an Agilent 6890 gas chromatograph (GC) coupled to an Agilent 5975 mass selective detector (MS) (Agilent Technologies, Waldbronn, Germany). The GC was equipped with a DB-5 capillary column (0.25mm ID × 30m; film thickness 0.25μm, J&W Scientific, Folsom, CA, USA). Helium was used as the carrier gas with a constant flow of 1mL/min. A temperature program from 60 °C to 300 °C with a 5 °C /min ramp and a final 10min at 300 °C was employed. Mass spectra were recorded in the EI mode with an ionization voltage of 70 eV and a source temperature of 230 °C. The software MassHunter Qualitative Analysis Navigator v.B.08.00 (Agilent Technologies) was used for data analysis. Identification of the compounds was accomplished by comparison of library data (NIST 20) with mass spectral data of commercially purchased standards for n-alkanes, diagnostic ions and retention indices. To calculate the relative abundance of CHC compounds, the area under each hydrocarbon peak in the chromatogram was quantified through integration and divided by the total area under all hydrocarbon peaks. Seventeen CHC compounds were found on the cuticle of *O. biroi*. These compounds are listed in **Supplementary Table S1**. To quantify the absolute abundance of CHC compounds of 1-day-old, 1-week-old, 1-month-old and 3-months-old ants (**Supplementary Figure S4A**), we added 10μL of C18 (10ng/μL) as an internal standard into each sample (amounting to 100ng of C18 per sample). C18 is not part of the ants’ natural CHC profile.

### Statistical analysis

All graphs were generated using GraphPad Prism 10.0.0. Statistical analyses were performed using R 4.3.2 (The R Foundation for Statistical Computing). Residuals were visually inspected for normality, and homogeneity of variances was evaluated with Levene’s tests. Generalized linear models (GLM) with a Poisson distribution, corrected for overdispersion, were used to analyze multiple comparisons of the total number of aggression events across focal genotypes, with host colony included as a fixed factor. When date was a significant random factor, generalized linear mixed models (GLMMs) with a negative binomial distribution were used. Kruskal-Wallis tests followed by post hoc comparisons with Bonferroni correction and ANOVA tests followed by Tukey’s post hoc tests were used to compare the total number of aggressions toward focal ants and the relative abundance of CHC compounds across more than two groups, when no significant effect of other factors (i.e., host colony and date) was detected. The Wilcoxon-Mann-Whitney test was used for pairwise comparisons of the total number of aggressions toward focal ants between wild-type and Bˈ ants. A linear regression was performed to assess the correlation between the number of host ant aggressions and the number of focal ant aggressions, using GraphPad Prism 10.0.0.

CHC profiles were visualized with non-metric multidimensional scaling (NMDS) based on Bray-Curtis dissimilarity matrices, implemented with the *metaMDS* function from the *vegan* package^65^. A discriminant analysis of principal components (DAPC) was performed using the *adegenet* package^66^ to visualize genotype-specific clustering of CHC profiles and temporal changes in CHC profiles of Bˈ ants raised in foster colonies over time. The effects of genotype and age on CHC profile composition were analyzed using permutational multivariate analysis of variance (ADONIS) on the same Bray-Curtis dissimilarity matrices, with genotype included as a fixed effect, using the *adonis* function from *vegan*^65^. Pairwise comparisons were then conducted using the *pairwiseAdonis* package (GitHub^67^), with p-values adjusted for multiple testing using the Benjamini-Hochberg (FDR) method.

## ACKNOWLEDGMENTS

We thank Yohann Chemtob for assisting with the video recording system and Tomas Kay for assistance with ant painting and helpful suggestions on the structure and writing of the manuscript. This work was supported by the European Molecular Biology Organization (EMBO) under award number ALTF 131-2022, the Leon Levy Foundation through the Leon Levy fellowship in Neuroscience, and the Price Family Center for the Social Brain through a Price fellowship (all to T.P.M.B.). M.R. was supported by an EMBO award (ALTF 955-2022) and a Human Frontier Science Program award (LT0023/2023-L). E.T.F. was supported by the Emmy Noether Program of the German Research Foundation (grant number: 511474012). T.S. was supported by funds from the University of Würzburg. Additional support was provided by the Howard Hughes Medical Institute, where D.J.C.K. is an investigator. This is Clonal Raider Ant Project paper number 42.

## AUTHOR CONTRIBUTIONS

T.P.M.B., M.R. and D.J.C.K. designed the study. T.P.M.B. and S.V.R. maintained and prepared ants for behavioral experiments. T.P.M.B. conducted behavioral experiments and analyses. T.P.M.B., M.R., E.T.F., and T.S. performed chemical experiments and analyses. T.P.M.B. and D.J.C.K. wrote the paper with help from M.R. and E.T.F., and T.P.M.B. prepared figures with input from D.J.C.K. D.J.C.K. supervised the project. All authors discussed the results and approved the final manuscript.

