## Supplemental material for "Tolerance toward foreigners in ants requires chronic exposure for establishment but only sporadic exposure for maintenance"

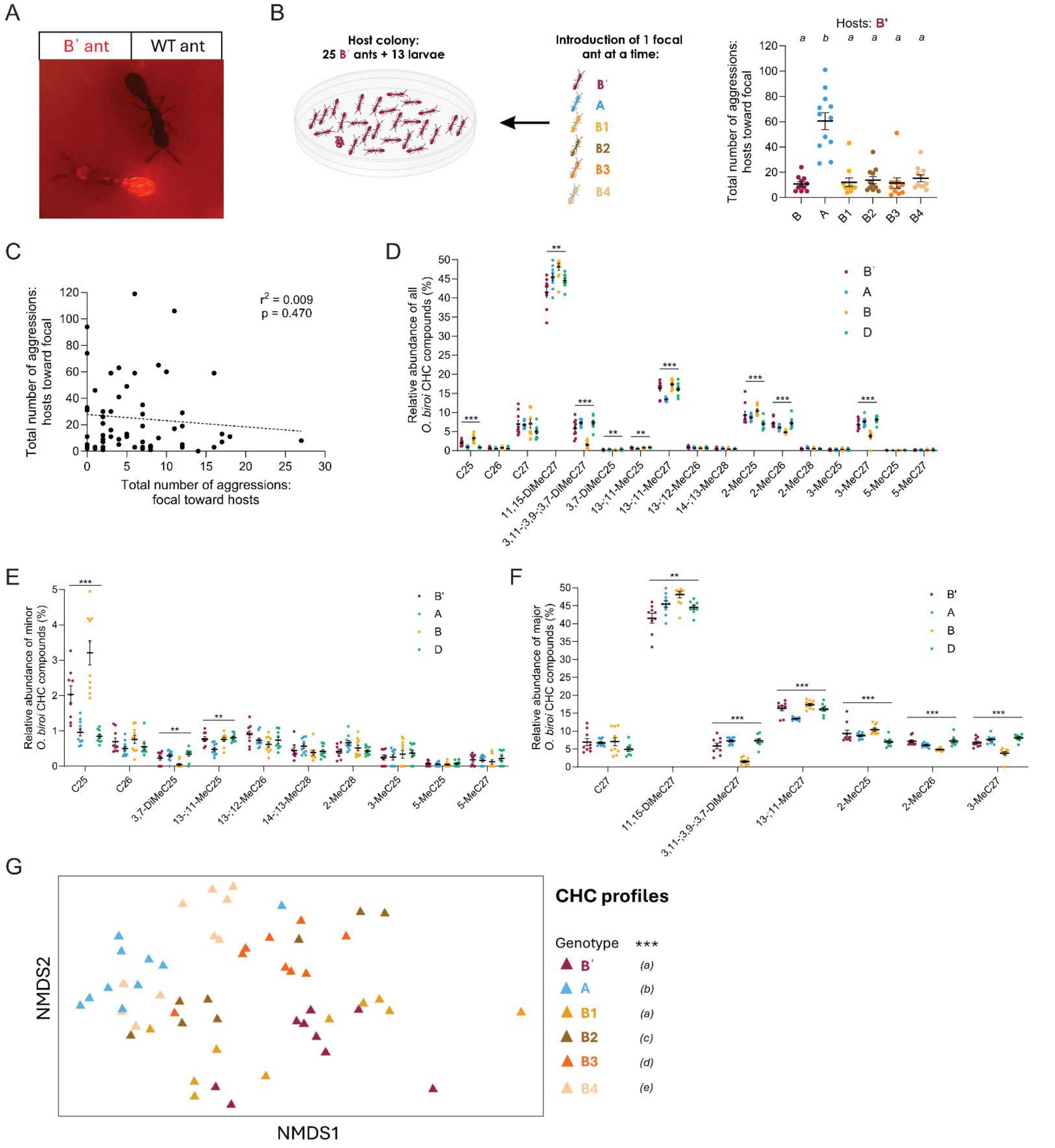


**Figure S1: Aggression toward foreign genotypes is stable over time.**

(A) Image of a B' ant and a wild-type (WT) ant acquired with an epifluorescence microscope. B' ants broadly express the red fluorescent protein dsRed, which allows us to easily distinguish and separate them from wild-type ants.

(B) Schematic and results of the behavioral assay. Focal ants of genotype B but from different stock colonies (B1-B4) were introduced one at a time to a host colony composed of 25 B' ants and 13 larvae. Focal ants from genotypes B' and A were also tested as negative and positive controls, respectively. Instances of aggressive

behavior were counted during 30 minutes. N=11-12 replicates per genotype. Error bars: standard error of the mean (SEM). Different letters indicate differences between focal genotypes ($p < 0.05$).
(C) Correlation between the number of aggressive behaviors exhibited by host ants toward focal ants and the number of aggressive behaviors exhibited by focal ants toward host ants. The $r^2$ and p values refer to the fit of a linear regression.
(D) Relative abundance of each of the 24 hydrocarbon compounds present on the *O. biroi* cuticle for genotypes B', A, B and D. Data points represent individual ants (N=9-10 replicates per genotype). Asterisks indicate differences between genotypes for each compound (*** $p < 0.001$, ** $p < 0.01$). Compounds that could not be consistently separated across experiments were lumped for this analysis and appear separated by semicolons.
Relative abundance of minor (E) and major (F) hydrocarbon compounds on the *O. biroi* cuticle were plotted separately to allow better visualization of the differences between genotypes. Asterisks indicate differences between genotypes for each compound (*** $p < 0.001$, ** $p < 0.01$).
(G) CHC profiles of B' (magenta) and A (blue) ants, as well as B ants from different laboratory stock colonies (different shades of yellow, orange and brown), visualized with an NMDS analysis. Data points represent individual ants (N=9-10 replicates per genotype). The NMDS analysis was conducted based on the relative abundance of each CHC compound across the different ant genotypes (a detailed list of the 24 compounds present in *O. biroi* in **Supplementary Table S1**). *** indicates an overall significant difference among genotypes ($p < 0.001$), and different letters indicate CHCs profile differences (multiple comparisons) between genotypes.
Details of statistical analyses are provided in **Supplementary Table S2.**

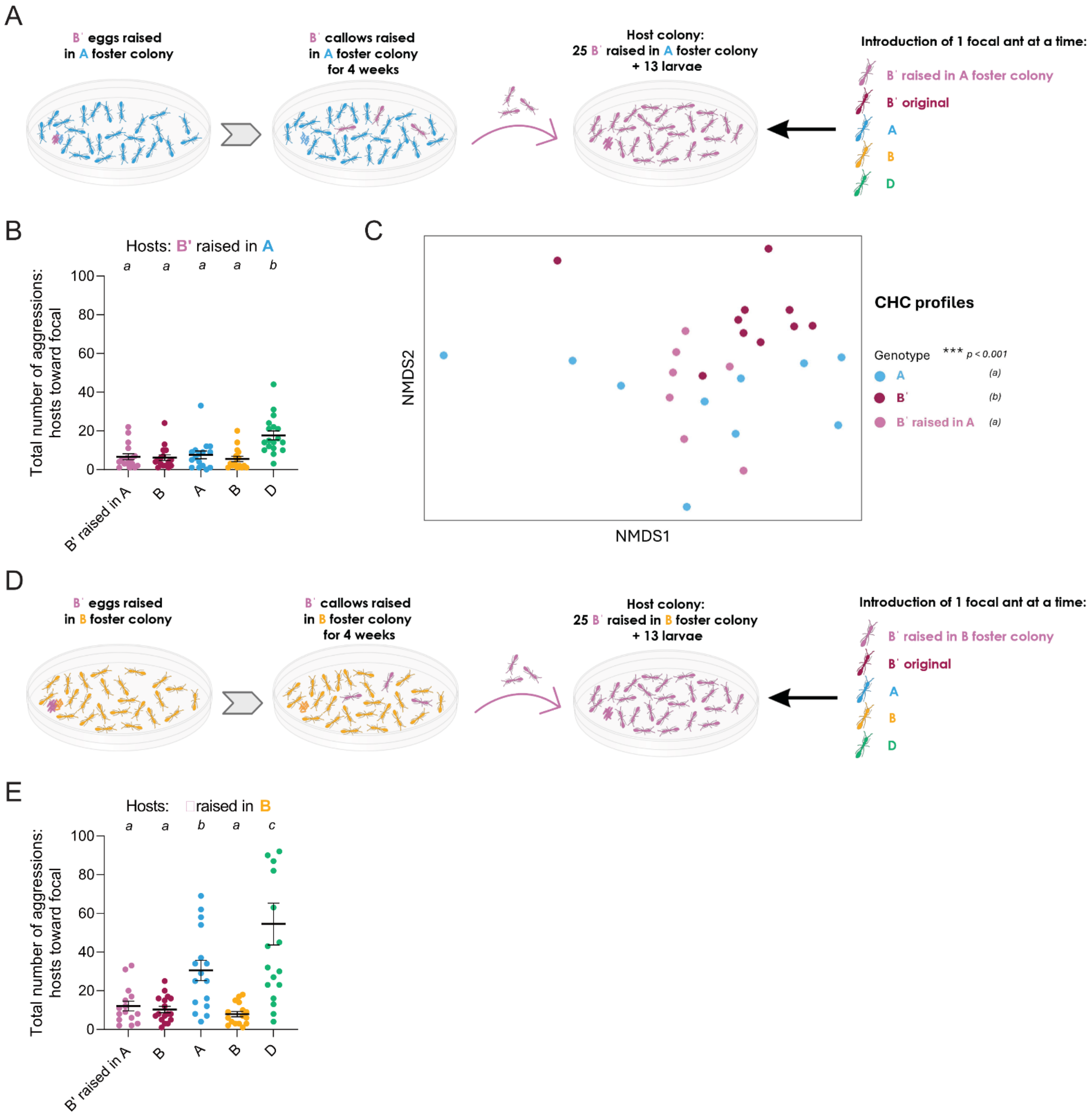


**Figure S2: Prolonged exposure establishes tolerance toward foreign genotypes.**

(A) Schematic of the behavioral assay. B' eggs were placed in genotype A foster colonies in a ratio of 1 B' egg to 8 foster ants. Once these B' eggs had developed into callow adults, these ants remained in the foster colonies for an additional four weeks. B' ants were then grouped into new colonies of 25 ants each, together with 13 larvae. Focal ants from different genotypes (B' raised in A, B' original, A, B and D) were then introduced to these B' host colonies one ant at a time, and aggression was recorded for 30 minutes.

(B) The total number of aggressive behaviors exhibited by B' ants raised in A foster colonies toward focal ants of different genotype. N=16-18 replicates per genotype. Error bars: standard error of the mean (SEM). Different letters indicate differences between focal genotypes ($p < 0.05$).

(C) CHC profiles of B' ants raised in A foster colonies (pink), original B' ants (magenta) and A ants (blue) visualized with an NMDS analysis. Data points represent individual ants (N=9-10 replicates per genotype). The NMDS analysis was conducted based on the relative abundance of each CHC compound across the different groups. *** indicates an overall significant difference among genotypes ($p < 0.001$), and different letters indicate CHC profile differences between genotypes after correcting for multiple comparisons.
(D) Schematic of the behavioral assay. Same as in (A), but with foster colonies of clonal line B.
(E) Total number of aggressive behaviors exhibited by B' ants raised in B foster colonies toward focal ants of different genotypes. N=15-18 replicates per genotype. Error bars: standard error of the mean (SEM). Different letters indicate differences between focal genotypes ($p < 0.05$).
Details of statistical analyses are provided in **Supplementary Table S2.**

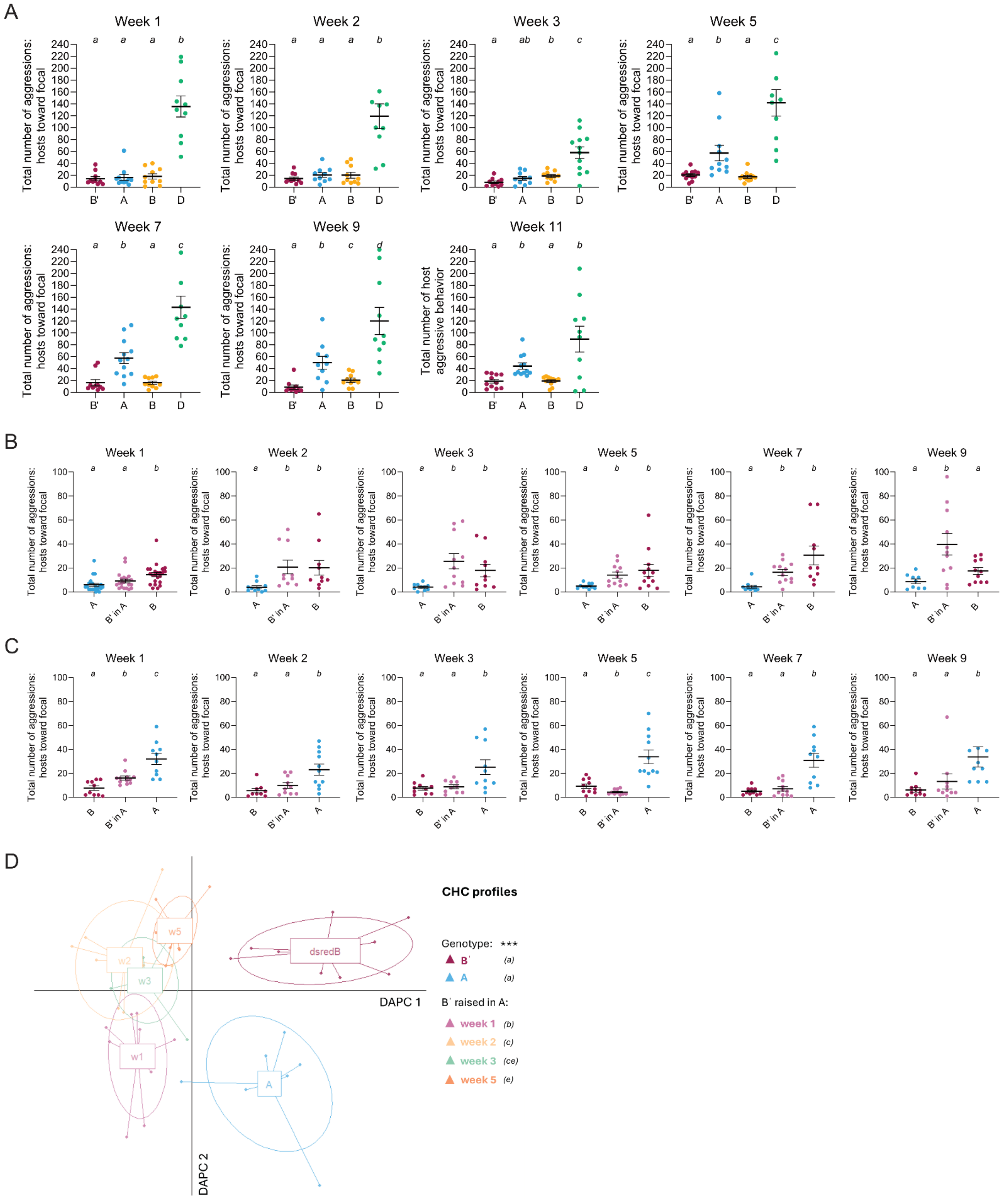


**Figure S3: Tolerance degrades with prolonged separation.**

(A) Total number of aggressive behaviors exhibited by B' ants raised in A foster colonies toward focal ants of genotypes B', A, B and D over time. N=10-12 replicates per genotype. Error bars: standard error of the mean (SEM). Different letters indicate differences between focal genotypes ($p < 0.05$).

(B) Total number of aggressive behaviors exhibited by A ants toward focal ants of genotypes B', B' raised in A and A over time. B' ants raised in A were used as focal ants 1, 2, 3, 5, 7 and 9 weeks after separation and isolation from A foster colonies. N=10-23 replicates per genotype. Error bars: standard error of the mean (SEM). Different letters indicate differences between focal genotypes ($p < 0.05$).

(C) Total number of aggressive behaviors exhibited by B' ants (original genotype) toward focal ants of genotypes B', B' raised in A and A over time. B' ants raised in A were used as focal ants 1, 2, 3, 5, 7 and 9 weeks after separation and isolation from A foster colonies. N=9-12 replicates per genotype. Error bars: standard error of the mean (SEM). Different letters indicate differences between focal genotypes ($p < 0.05$).

(D) CHC profiles of genotypes A (blue), B' original (magenta), and B' raised in A colonies at 1, 2, 3 and 5 weeks after separation from their foster colonies (pink, yellow, green, orange), visualized using a discriminant analysis of principal components (DAPC). Data points represent individual ants (N=9-10 replicates per condition). The DAPC was performed on the relative abundance of 24 CHC compounds identified in *O. biroi* (see **Supplementary Table S1**). *** indicates an overall significant difference among genotypes ($p < 0.001$), and different letters denote significant pairwise differences in CHC profiles between groups ($p < 0.05$).

Details of statistical analyses are provided in **Supplementary Table S2.**

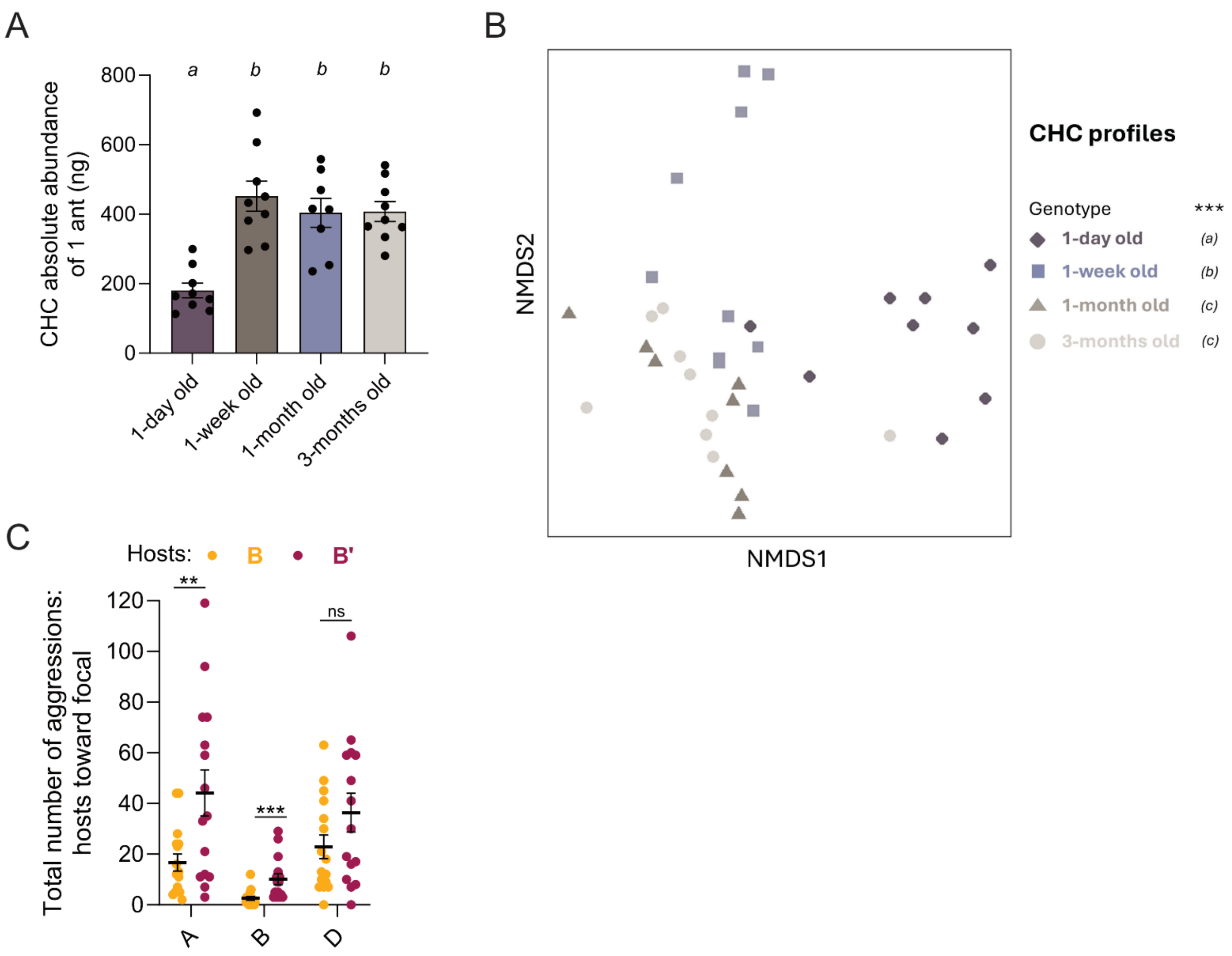


**Figure S4: Related to STAR methods section.**
(A) Quantification of the total amount of CHCs on the cuticle of 1-day, 1-week, 1-month and 3-months old ants from clonal line B. All ants came from the same stock colony. Analyses were calibrated using a C18 standard (10ng/µL). N=8-9 replicates per condition. Different letters indicate differences between age cohorts ($p < 0.05$).
(B) CHC profiles of genotype B ants of different ages, visualized with an NMDS analysis. Data points represent individual ants (N=9-10 replicates per age cohorts). The analysis was conducted based on the relative abundance of CHC compounds across the different age cohorts (a detailed list of the 24 compounds present in *O. biroi* is provided in **Supplementary Table S1**). *** indicates an overall significant difference among age cohorts ($p < 0.001$), and different letters indicate CHC profile differences between age cohorts after correcting for multiple comparisons.
(C) Total number of aggressive behaviors exhibited by genotype B (yellow) and B' (magenta) host ants toward focal ants of genotypes A, B and D. N=15-16 replicates per genotype. Error bars: standard error of the mean (SEM). Asterisks indicate differences between B and B' host colonies: ***$p < 0.001$, **$p < 0.01$, 'ns' not significant.
Details of statistical analyses are provided in **Supplementary Table S2.**

**Supplementary Video S1. Non-nestmate discrimination in the clonal raider ant.**

Examples of two aggressive behaviors, biting and bending the gaster, exhibited by clonal raider ants during non-nestmate discrimination.

**Supplementary Table S1:** The 24 CHC compounds identified in *O. biroi.* For each compound, we report the GC-MS retention index (RI) and the relative abundance (mean ± standard deviation) across the four ant clonal genotypes (B', A, B and D). Semicolons separate compounds that were detected but could not be reliably separated in all samples. These compounds were therefore lumped in our chemical analyses and are given as single entries in the table.

| **Compound** | **RI** | **B'** | **A** | **B** | **D** |
|---|---|---|---|---|---|
| C25 | 2,500 | 2.04 ± 0.24 | 0.70 ± 0.07 | 1.15 ± 0.08 | 0.35 ± 0.03 |
| 13-;11-MeC25 | 2,533 | 0.78 ± 0.06 | 0.55 ± 0.04 | 0.95 ± 0.05 | 0.71 ± 0.07 |
| 5-MeC25 | 2,549 | 0.07 ± 0.02 | 0.08 ± 0.02 | 0.09 ± 0.01 | 0.07 ± 0.02 |
| 3-;2-MeC25 | 2,563 | 9.60 ± 0.89 | 8.97 ± 0.44 | 11.18 ± 0.30 | 8.44 ± 0.18 |
| C26; 3,7-DiMeC25 | 2,600 | 0.92 ± 0.05 | 0.80 ± 0.04 | 0.62 ± 0.05 | 0.55 ± 0.07 |
| 13-;12-MeC26 | 2,632 | 0.90 ± 0.10 | 0.91 ± 0.07 | 0.97 ± 0.06 | 0.81 ± 0.10 |
| 3-;2-MeC26 | 2,663 | 7.18 ± 0.41 | 5.88 ± 0.19 | 5.04 ± 0.14 | 4.77 ± 0.15 |
| C27 | 2,700 | 6.94 ± 1.00 | 5.79 ± 0.36 | 4.56 ± 0.37 | 3.38 ± 0.19 |
| 13-;11-MeC27 | 2,730 | 16.51 ± 0.70 | 14.90 ± 0.43 | 18.68 ± 0.25 | 17.85 ± 0.18 |
| 11,15-DiMeC27 | 2,757 | 41.48 ± 1.33 | 45.54 ± 0.69 | 48.07 ± 1.03 | 46.15 ± 0.76 |
| 5-;3-MeC27 | 2,770 | 6.83 ± 0.42 | 6.90 ± 0.28 | 4.10 ± 0.17 | 7.34 ± 0.31 |
| 3,11-;3,9-;3,7-DiMeC27 | 2,801 | 5.87 ± 0.77 | 7.45 ± 0.25 | 3.56 ± 0.24 | 8.29 ± 0.43 |
| 14-;13-MeC28 | 2,827 | 0.44 ± 0.08 | 0.81 ± 0.16 | 0.26 ± 0.05 | 0.63 ± 0.11 |
| 2-MeC28 | 2,853 | 0.42 ± 0.06 | 0.71 ± 0.08 | 0.77 ± 0.04 | 0.66 ± 0.09 |

**Supplementary Table S2**: Summary of statistical analyses.

| Figure | Test | Response variable | Explanatory factor | Results | Post-hoc test (only p-values < 0.05) |
|---|---|---|---|---|---|
| 1C | Quasi-Poisson GLM | Total number of aggressions | Focal genotype [A, B, D, B'] | F(3,56) = 16.04, p < 0.001 | B vs A: p < 0.001; B vs D: p = 0.001; B' vs A: p < 0.001; B' vs D: p < 0.001 |
| 1D | Adonis test | CHC profile | Genotype [A, B, D, B'] | F(3,36) = 11.16, p < 0.001 | B' vs B: p < 0.001; B' vs D: p < 0.001; A vs B: p < 0.001; A vs D: p < 0.001; B vs D: p < 0.001; A vs B': p = 0.015 |
| 1E | Kruskal-Wallis test | Total number of aggressions | Focal genotype [A, B, D, B'] | Week 1: $\chi^2$ = 21.17, df = 3, p < 0.001; Week 2: $\chi^2$ = 21.14, df = 3, p < 0.001; Week 3: $\chi^2$ = 19.74, df = 3, p < 0.001; Week 5: $\chi^2$ = 25.45, df = 3, p < 0.001; Week 7: $\chi^2$ = 26.08, df = 3, p < 0.001; Week 9: $\chi^2$ = 27.72, df = 3, p < 0.001; Week 11: $\chi^2$ = 24.32, df = 3, p < 0.001 | Week 1: A vs B: p < 0.001; A vs D: 0.018; A vs B': p < 0.001; B vs D: p = 0.024.<br>Week 2: A vs B: p < 0.001; A vs B': p = 0.002; B vs D: p < 0.001; D vs B': p = 0.006.<br>Week 3: A vs B: p = 0.004; A vs B': p = 0.005; B vs D: p < 0.001; D vs B': p < 0.001.<br>Week 5: A vs B: p < 0.001; A vs D: p = 0.008; A vs B': p = 0.006; B vs D: p < 0.001; D vs B': p < 0.001.<br>Week 7: A vs B: p = 0.001; A vs D: p = 0.001; A vs B': p = 0.009; B vs D: p < 0.001; D vs B': p < 0.001.<br>Week 9: A vs B: p < 0.001; A vs D: p = 0.002; A vs B': p = 0.002; B vs D: p < 0.001; D vs B': p < 0.001.<br>Week 11: A vs B: p < 0.001; A vs B': p < 0.001; B vs D: p < 0.001; D vs B': p < 0.001. |
| 2B | Quasi-Poisson GLM | Total number of aggressions | Focal genotype [A, B, D, B', B' raised in B] | F(4,59) = 13.20, p < 0.001 | B vs A: p < 0.001; D vs A: p = 0.02; B' in B vs A: p < 0.001; B' vs A: p = 0.004; D vs B: p = 0.028; B' in B vs D: p = 0.044 |
| 2C | Quasi-Poisson GLM | Total number of aggressions | Focal genotype [A, B, D, B', B' raised in A] | F(4,59) = 6.21, p < 0.001 | D vs A: p = 0.002; D vs B: p = 0.002; B vs A: p < 0.001; D vs B': p = 0.002 |
| 2D | Negative Binomial GLMM | Total number of aggressions | Focal genotype [A, B, D, B', B' raised in D] | $\chi^2$ = 37.73, df = 4, p < 0.001 | B vs A: p < 0.001; D vs A: p < 0.001; B' vs A: p < 0.001; B' in D vs A: p < 0.001 |
| 2E | Adonis test | CHC profile | Original genotypes [B', A, B, D] | F(3,65) = 14.53, p < 0.001 | B' vs A: p = 0.007; B' vs B: p < 0.001; B' vs D: p < 0.001; A vs B: p < 0.001; A vs D: p < 0.001; B vs D: p < 0.001; |
| | | | WT genotypes [A, B, D] vs cross-fostered B' [B' raised in A, B and D] | F(1,67) = 1.61, p = 0.175 | B vs B' in B: p > 0.05<br>D vs B' in D: p > 0.05<br>A vs B' in A: p = 0.021 |
| 3B | Kruskal-Wallis test | Total number of aggressions toward A focal ants | Week [weeks 1, 2, 3, 5, 7, 9, 11] | $\chi^2$ = 36.81, df = 6, p < 0.001 | w1 vs w11: p < 0.001; w1 vs w5: p < 0.001; w1 vs w7: p < 0.001; w1 vs w9: p < 0.001; w11 vs w2: p = 0.006; w11 vs w3: p < 0.001; w2 vs w5: p = 0.005; w2 vs w7: p < 0.001; w2 vs w9: p = 0.017; w3 vs w5: p < 0.001; w3 vs w7: p < 0.001; w3 vs w9: p < 0.001 |
| 3C | Kruskal-Wallis test | Total number of aggressions | Focal genotype [A, B', B' raised in A] | Week 1: $\chi^2$ = 16.52, df = 2, p < 0.001; Week 2: $\chi^2$ = 12.62, df = 2, p = 0.002; Week 3: $\chi^2$ = 11.32, df = 2, p = 0.003 | Week 1: A vs B': p < 0.001; B' vs B' in A: p = 0.015. Week 2: A vs B': p = 0.001; A vs B' in A: p < 0.001<br>Week 3: A vs B': p = 0.013; A vs B' in A: p < 0.001 |

| | | | | | |
|---|---|---|---|---|---|
| 3D | Kruskal-Wallis test | Total number of aggressions | Focal genotype [A, B', B' raised in A] | Week 1: $\chi^2 = 18.52$, df = 2, $p < 0.001$; Week 2: $\chi^2 = 10.59$, df = 2, p = 0.005; Week 3: $\chi^2 = 7.71$, df = 2, p = 0.021 | Week 1: A vs B': p < 0.001; A vs B' in A: p = 0.002; B' vs B' in A: p = 0.003<br>Week 2: A vs B': p < 0.001; A vs B' in A: p = 0.03<br>Week 3: A vs B': p = 0.009; A vs B' in A: p = 0.043 |
| 4B | Wilcoxon-Mann-Whitney test | Total number of aggressions for each experimental week | Focal genotype [A, B'] | Week 1: W= 98, p = 0.809<br>Week 2: W=110, p = 0.597<br>Week 3: W = 107, p = 0.419<br>Week 4: W = 94, p = 0.903<br>Week 5: W = 64.5, p = 0.951<br>Week 7: W = 128.5, p = 0.006 | NA |
| S1B | Quasi-Poisson GLM | Total number of aggressions | Focal genotype [A, B', B1, B2, B3, B4] | $F(5,61) = 11.36$, $p < 0.001$ | A vs B1: p < 0.001; A vs B2: p < 0.001; A vs B3: p < 0.001; A vs B4: p < 0.001; A vs B': p < 0.001 |
| S1C | Linear correlation | Number of host aggressions | Number of focal aggressions | F = 0.528, $R^2 = 0.009$, p = 0.470 | NA |
| S1D-F | Kruskal-Wallis test | Relative abundance of each CHC compound | Genotype [A, B, D, B'] | 2-MeC25: $\chi^2 = 18.83$, df = 3, p < 0.001; 2-MeC26: $\chi^2 = 25.18$, df = 3, p < 0.001; 2-MeC28: $\chi^2 = 7.04$, df = 3, p = 0.07; 3-MeC25: $\chi^2 = 1.75$, df = 3, p = 0.62; 3-MeC27: $\chi^2 = 24.21$, df = 3, p < 0.001; 5-MeC25: $\chi^2 = 1.07$, df = 3, p = 0.78; 5-MeC27: $\chi^2 = 1.92$, df = 3, p = 0.59; 3,7-DiMeC25: $\chi^2 = 14.86$, df = 3, p = 0.002; 11,15-DiMeC27: $\chi^2 = 15.42$, df = 3, p < 0.001; 13-;11-MeC25: $\chi^2 = 15.68$, df = 3, p < 0.001; 13-;11-MeC27: $\chi^2 = 18.09$, df = 3, p < 0.001; 13-; 12-MeC26: $\chi^2 = 4.9$, df = 3, p = 0.18; 14-;13-MeC28: $\chi^2 = 4.93$, df = 3, p = 0.18; 3,11-;3,9-;3,7-DiMeC27: $\chi^2 = 22.64$, df = 3, p < 0.001; C25: $\chi^2 = 27.34$, df = 3, p < 0.001; C26: $\chi^2 = 4.76$, df = 3, p = 0.19; C27: $\chi^2 = 5.16$, df = 3, p = 0.16 | 2-MeC25: A vs B: p = 0.035; A vs D: p = 0.006; B vs D: p < 0.001; B vs B': p = 0.023; D vs B': p = 0.014. 2-MeC26: A vs B: p < 0.001; A vs D: p = 0.0075; A vs B': p = 0.018; B vs D: p < 0.001; B vs B': p < 0.001. 3-MeC27: A vs B: p < 0.001; B vs D: p < 0.001; B vs B': p < 0.001; D vs B': p = 0.035. 3,7-DiMeC25: A vs B; p = 0.003; B vs D: p < 0.001; B vs B': p = 0.07. 11,15-DiMeC27: A vs B': p = 0.031; B vs D: p = 0.017; B vs B': p < 0.001. 13-;11-MeC25: A vs B: p < 0.001; A vs D: p < 0.001; A vs B': p < 0.001. 13-;11-MeC27: A vs B: p < 0.001; A vs D: p = 0.005; A vs B': p = 0.001. 3,11-;3,9-;3,7-DiMeC27: A vs B: p < 0.001; B vs D: p < 0.001; B vs B': p < 0.001. C25: A vs B: p < 0.001; A vs B': p < 0.001; B vs D: p < 0.001; D vs B': p< 0.001 |
| S1G | Adonis test | CHC profile | Genotype [A, B', B1, B2, B3, B4] | $F(5,50) = 11.31$, $p < 0.001$ | All comparisons p < 0.001 except: B1 vs B3: p = 0.01; B2 vs A: p = 0.008; B' vs B1: p = 0.268 |
| S2B | Quasi-Poisson GLM | Total number of aggressions | Focal genotype [A, B, D, B', B' raised in A] | $F(4,78) = 7.10$, $p < 0.001$ | D vs A: p = 0.012; D vs B: p < 0.001; B' in A vs D: p = 0.002; B' vs D: p = 0.002 |
| S2C | Adonis test | CHC profile | Genotype [A, B', B' raised in A] | $F(2,24) = 3.86$, p = 0.005 | B' in A vs B': p = 0.019; A vs B': p = 0.012 |
| S2E | Quasi-Poisson GLM | Total number of aggressions | Focal genotype [A, B, D, B', B' raised in B] | $F(4,77) = 23.59$, $p < 0.001$ | B vs A: p < 0.001; D vs A: p = 0.016; B' in B vs A: p = 0.016; B' vs A: p = 0.002; B vs D: p < 0.001; B' in B vs D: p < 0.001; B' vs D: p < 0.001 |
| S3A | Kruskal-Wallis test | Total number of aggressions | Focal genotype [A, B, D, B'] | Week 1: $\chi^2 = 22.14$, df = 3, p < 0.001; Week 2: $\chi^2 = 20.69$, df = 3, p < 0.001; Week 3: $\chi^2 = 19.51$, df = 3, p < 0.001; Week 5: $\chi^2 = 31.66$, df = 3, p < 0.001; | Week 1: A vs D: p < 0.001; B vs D: p < 0.001; D vs B': p < 0.001.<br>Week 2: A vs D: p < 0.001; B vs D: p < 0.001; D vs B': p < 0.001 |

| | | | | | |
|---|---|---|---|---|---|
| | | | | Week 7: $\chi^2 = 31.13$, df = 3, $p < 0.001$; Week 9: $\chi^2 = 26.38$, df = 3, $p < 0.001$; Week 11: $\chi^2 = 18.53$, df = 3, $p < 0.001$ | Week 3: A vs D: $p < 0.001$; B vs D: $p = 0.022$; B vs B': $p = 0.032$; D vs B': $p < 0.001$<br>Week 5: A vs B: $p < 0.001$; A vs D: $p < 0.001$; A vs B': $p < 0.001$; B vs D: $p < 0.001$; D vs B': $p < 0.001$<br>Week 7: A vs B: $p < 0.001$; A vs D: $p < 0.001$; A vs B': $p < 0.001$; B vs D: $p < 0.001$; D vs B': $p < 0.001$<br>Week 9: A vs B: $p = 0.046$; A vs D: $p = 0.013$; A vs B': $p < 0.001$; B vs D: $p < 0.001$; B vs B': $p = 0.031$; D vs B': $p < 0.001$<br>Week 11: A vs B: $p < 0.001$; A vs B': $p = 0.001$; B vs D: $p = 0.001$; D vs B': $p = 0.003$ |
| S3B | Kruskal-Wallis test | Total number of aggressions | Focal genotype [A, B', B' raised in A] | Weeks 1&2&3: see 4D; Week 5: $\chi^2 = 10.78$, df = 2, $p < 0.001$; Week 7: $\chi^2 = 15.42$, df = 2, $p < 0.001$; Week 9: $\chi^2 = 9.92$, df = 2, $p = 0.01$ | Weeks 1&2&3: see 4D.<br>Week 5: A vs B': $p = 0.003$; A vs B' in A: $p = 0.002$<br>Week 7: A vs B': $p < 0.001$; A vs B' in A: $p < 0.001$<br>Week 9: A vs B' in A: $p = 0.001$ |
| S3C | Kruskal-Wallis test | Total number of aggressions | Focal genotype [A, B', B' raised in A] | Weeks 1&2&3: see 4E; Week 5: $\chi^2 = 21.14$, df = 2, $p < 0.001$; Week 7: $\chi^2 = 15.12$, df = 2, $p < 0.001$; Week 9: $\chi^2 = 14.83$, df = 2, $p < 0.001$ | Weeks 1&2&3: see 4E.<br>Week 5: A vs B': $p < 0.001$; A vs B' in A: $p < 0.001$; B' vs B' in A: $p = 0.019$<br>Week 7: A vs B': $p < 0.001$; A vs B' in A: $p < 0.001$<br>Week 9: A vs B': $p < 0.001$; A vs B' in A: $p < 0.001$ |
| S3D | Adonis test | CHC profile | Genotype [A, B', B' raised in A] and week [weeks 1, 2, 3, 5] | $F(5,48) = 10.26$, $p < 0.001$ | All comparisons $p < 0.05$ except: w2 vs w3: $p = 0.107$, w3 vs w5: $p = 0.134$, B' vs A: $p = 0.137$ |
| S4A | Anova test | Total number of aggressions | Age [1-day, 1-week, 1-month, 3-months old] | $F(3,31) = 12.81$, $p < 0.001$ | 1day vs 1month: $p < 0.001$; 1day vs 1week: $p < 0.001$; 1day vs 3months: $p < 0.001$ |
| S4B | Adonis test | CHC profile | Age [1-day, 1-week, 1-month, 3-months old] | $F(3,32) = 10.91$, $p < 0.001$ | 1day vs 1month: $p < 0.001$; 1month vs 1week: $p < 0.001$; 1day vs 3months: $p < 0.001$; 3months vs 1week: $p < 0.001$; 1day vs 1week: $p < 0.001$ |
| S4C | Wilcoxon-Mann-Whitney test | Total number of aggressions toward A | Host genotype [B, B'] | W = 177, $p = 0.025$ | NA |
| | | Total number of aggressions toward B | | W = 208, $p < 0.001$ | NA |
| | | Total number of aggressions toward D | | W = 152, $p = 0.212$ | NA |
